# Genome Tree of Life: *Deep Burst* of Organism Diversity

**DOI:** 10.1101/756155

**Authors:** JaeJin Choi, Sung-Hou Kim

## Abstract

An organism Tree of Life (*organism ToL*) is a conceptual and metaphorical tree to capture a simplified narrative of the evolutionary course and kinship among the extant organisms of today. Such tree cannot be experimentally validated but may be reconstructed based on characteristics associated with the extant organisms. Since the whole genome sequence of an organism is, at present, the most comprehensive *descriptor* of the organism, a *genome Tol* can be an empirically derivable surrogate for the organism ToL. However, a genome ToL has been impossible to construct because of the practical reasons that experimentally determining the whole genome sequences of a large number of diverse organisms was technically impossible. Thus, for several decades, *gene ToLs*, based on *selected genes*, have been commonly used as a surrogate for the organisms ToL. This situation changed dramatically during the last several decades due to rapid advances in DNA sequencing technology. Here we describe the main features of a genome ToL that are different from those of the broadly accepted gene ToLs: (a) the first two organism groups to emerge are the founders of prokarya and eukarya, (b) they diversify into six large groups and all the *founders* of the groups have *emerged* in a *“Deep Burst”* at the very beginning period of the emergence of Life on Earth and (c) other differences are notable in the order of emergence of smaller groups.

**Significance Statement:** *Tree of Life* is a conceptual and metaphorical tree that captures a simplified narrative of the *evolutionary course* and *kinship* among all living organisms of today. Since the whole genome sequence information of an organism is, at present, the most comprehensive description of the organism, we reconstructed a *Genome Tree of Life* using the proteome information from the whole genomes of over 4000 different living organisms on Earth. It suggests that (a) the first two primitive organism groups to emerge are the founders of prokarya and eukarya, (b) they diversify into six large groups, and (c) all the *founders* of the groups have *emerged* in a *“Deep Burst”* at the very beginning period of the emergence of Life on Earth.

## Introduction

### Organism Phylogeny vs. Gene Phylogeny

For decades, due to the technical difficulties of whole genome sequencing, the *gene Tree of Life* (ToL), constructed based on the information of a set of *selected genes*, has been used most commonly as a surrogate for the *organism ToL* despite the fact that gene ToLs most likely infer the evolutionary relationship of only the selected genes, not of the *organisms*. Furthermore, there are various other intrinsic limitations and confounding issues associated with the construction and interpretation of gene ToLs (1). Many gene ToLs have been constructed based on different sets of the selected genes, new or increased number of extant organisms, and other inputs combined with various different gene-based analysis methods (1-8). They showed mostly good agreements on the clading of organism groups, but with varying degrees of disagreements on the branching orders and branching time of the groups, especially at deep tree branching levels. Thus, it became increasingly uncertain (1, 9,10) whether *gene* ToLs are appropriate surrogates for the organism ToL. In addition, an important issue of rooting ToLs has not been well resolved and still been debated (11).

These and other issues of gene ToLs highlight the need for an alternative surrogate for the organism ToL built based on as *completely different assumptions* as possible from those of gene ToLs. A genome ToL based on *Information Theory* (12) may provide an independent and alternative view of the organism ToL.

### Genome Tree of Life

The whole genome sequence information of an extant organism can be considered, at present, as the most comprehensive *digital information* of the organism for its survival and reproduction in its current environment and ecology. How to format such information system (“descriptor”) and quantitatively compare a pair of the systems (“distance measure” for the degree of difference) are well developed in Information Theory, not only for digitally encoded electronic signals and images, but also for natural language systems, such as books and documents (13).

#### “Descriptor”

In this study we use the descriptor of Feature Frequency Profile (FFP) method (14; see FFP method in Materials and Methods) to describe the *whole genome sequence information* of an organism, be it the DNA sequence of the whole genome, the RNA sequence of the transcriptome or the amino acid sequence of the proteome of the organism. Briefly, a Feature in FFP is an adaptation of “n-grams” or “k-mers” used to describe a sentence, a paragraph, a chapter, or a whole book (15) in Information Theory and Computational Linguistics, where an n-gram is a string of n “charactors” (all alphabets plus space and delimiter such as comma, period etc.). But, for FFP method we treat the whole genome information of an organism as a book of alphabets without spaces and delimiters, and represent the book by its FFP, which is the collection of all *unique* n-grams and their frequencies in the book. It is important to note that such FFP of an organism has all the information necessary to reconstruct the original genome sequence information of the organism. The “optimal n” of the n-gram, the most critical parameter, for the construction of the genome ToL can be empirically obtained under a given criterion (see Choice of the “best” Descriptor and the “optimal” Feature length in Materials and Methods). For the criterion of the most topologically stable ToL, it usually ranges, depending on the size of the sample and the types of information (“alphabets” of genome information), between 10 – 15 for proteome sequences and 20 – 26 for genome or transcriptome sequences. For this study, we found that the proteome sequence with the optimal Feature length of 12 amino acids or longer yields the most topologically stable genome ToL (see Supplemental Fig. S1.)

#### “Distance measure”

As for the measure to estimate the degree of difference between two FFPs we use the *genomic divergence* as calculated by Jensen-Shannon Divergence (16) (for details see Jensen-Shannon Divergence and cumulative branch-length in Materials and Methods), an information-theoretical function for estimating the degree of divergence between two linear information systems. For this study, the divergence is assumed to be caused by the *changes* in genome sequence by all known and unknown mechanisms. Such divergences for all pairs of organisms, then, can be used to construct the “divergence distance matrix” needed to build a genome ToL.

#### Outgroup

The descriptor and the distance measure used in the FFP method provide another important advantage for constructing the genome sequence of an “artificial organism” that may be used as an outgroup member of a ToL. An artificial genome can be constructed to have the same genome size and composition of genomic alphabets as one of the real genomes of an extant organism in the study population, but the sequence of the alphabets are shuffled within the real genome, such that it has no information of sustaining and reproducing Life. Since the FFP method is one of the “alignment-free” methods which do not depend on the multiple sequence alignment of long stretches of sequences common among *all members* of the study population, such artificial organisms have been used successfully as members of an out-group in constructing the rooted whole genome trees for prokaryotes (17) and fungi (18).

### Pool of “founding ancestors” vs. common ancestor

Recent observations prompted us to revisit the meaning of *internal nodes* for this study: Whole genome sequences of a very large members of *Homo sapiens* and *E. coli* species have been experimentally determined in the last decade. They revealed that the extent of the genomic divergence and variation among the members in a species is very broad even after a short period of the evolutionary time of the species (19, 20,21) and even under a constant environment in the case of *E. coli* (21). Thus, an internal node can be considered, in this study, as a *pool of founding ancestors* (FA) with *wide genomic diversity*, from which the founders (small subpopulation of precursors) for the new groups emerge, or “sampled/selected”, under, for example, drastic changes in local environment and ecology. This “mosaic” feature of the internal node is conceptually different from the “clonal” feature of the node as a common ancestor, from which two or more descendants with *high genomic similarity* branch out (see Supplemental Fig. S2).

### Objective

Early on we experimented, optimized and tested the applicability of this Information Theory–based FFP method for constructing the rooted genome trees of prokaryotes (17) and fungi (18). Here we present a rooted genome ToL for all organisms for which the information of complete or near-complete genome sequence are publicly available (4023 “species” as of initiation of this study). It is hoped that the genome ToL adds a fundamentally new viewpoint to those of the existing gene ToLs and stimulates improvement of current methods as well as development of additional new methods in expectation of a much larger scale ToLs to cover new organisms of broader diversity in as yet unexplored environments on the Earth’s surface as well as in vast marine and subterranean worlds not yet surveyed.

## Results

In this section, we present our observations of the features in our genome ToL. Associated implications and narratives will be presented in Discussions section. To highlight the similarities and differences of the features of the genome ToL from those of gene ToLs, we present our results from two viewpoints: (a) identification of large groups and the *topological relationship* among the groups showing the order of emergence and (b) relative magnitude in *cumulative branch lengths* among the founders of all the groups to estimate the relative extent of *evolutionary progression* toward the extant organisms of each founder of the groups.

Since the grouping and the branching order of the groups in the genome ToL do not always agree with those in the gene ToLs, we use the following descriptions for the groups at various branching levels: “Supergroup”, “Majorgroup”, “Group”, and “Subgroup”. The generic labels of the groups in the genome ToL are assigned and the corresponding taxon names from National Center for Biotechnology Information (NCBI; see Materials and Methods), which are mostly based on the clading pattern in gene trees and characteristic phenotypes at the time of naming, are also listed for comparison.

### A. Grouping of extant organisms: Two Supergroups, six Majorgroups, and 35+ Groups

Figure 1 shows that, at the deepest level, two Supergroups emerge as indicated by the two-colored inner circular band: Red colored portion corresponds to Prokarya (“Akarya” (22) may be a more appropriate name, because the founders of Prokarya do not emerge before the founders of Eukarya in our ToL (see Fig. 2)) and the blue to Eukarya. At the next level, six Majorgroups emerge as indicated in the outer circular band by six colors. Finally, thirty-five Groups emerge as indicated by the small circles with their genome ToL labels (next to the circles) and corresponding scientific or common names used in NCBI database (outside of the circular bands).

**Fig. 1:**
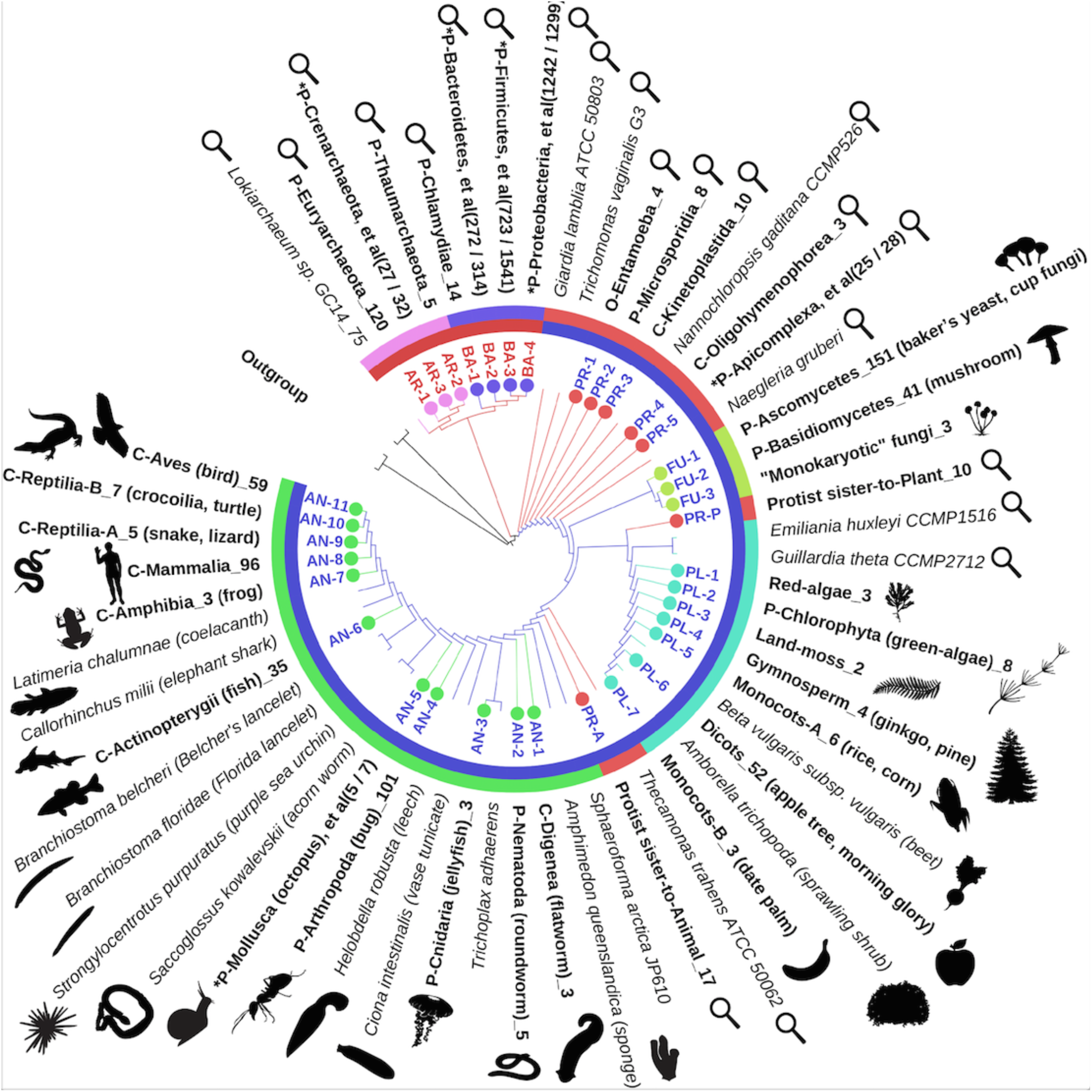
Topology and Branching order of the genome ToL. A circos ITOL (42) diagram highlighting the topology and branching order of the genome ToL of all collapsed clades (small colored circles) as well as most of “singletons” (in italics) in the study population. The branch-lengths are ignored. The inner ring has two colors for two Supergroups: Red corresponds to Prokarya (or Akarya) and blue to Eukarya. Outer ring has six colors for six Majorgroups: pink for AR (Archaea), blue for BA (Bacteria), orange for PR (Protists in three types: PR-1~5, PR-P, and PR-A), pale green for FU (Fungi), cyan for PL(Plants) and green for AN(Animals). The labels for 35 Groups (3 for AR, 4 for BA, 7 for PR in three Groups, 3 for FU, 7 for PL, and 11 for AN) are shown next to the small circles, and their corresponding scientific names according to NCBI taxonomy (preceded by taxonomic level of Phylum (P), Class(C), or Order (O)) are shown outside of the outer ring followed by the number of samples in this study. Common names and silhouettes of one or more examples are also shown. The symbol of a magnifying glass represents microbial organisms not visible by human eyes. Asterisks are for the clades with “mixed” names: For example, the group “*p-Bacteroidetes (272/314)” consists of 314 members of several bacterial phyla, of which 272 are Bactroidetes, the phylum with the largest sample size in the group. The visualization of ToL was made using ITOL(42).

**Fig. 2:**
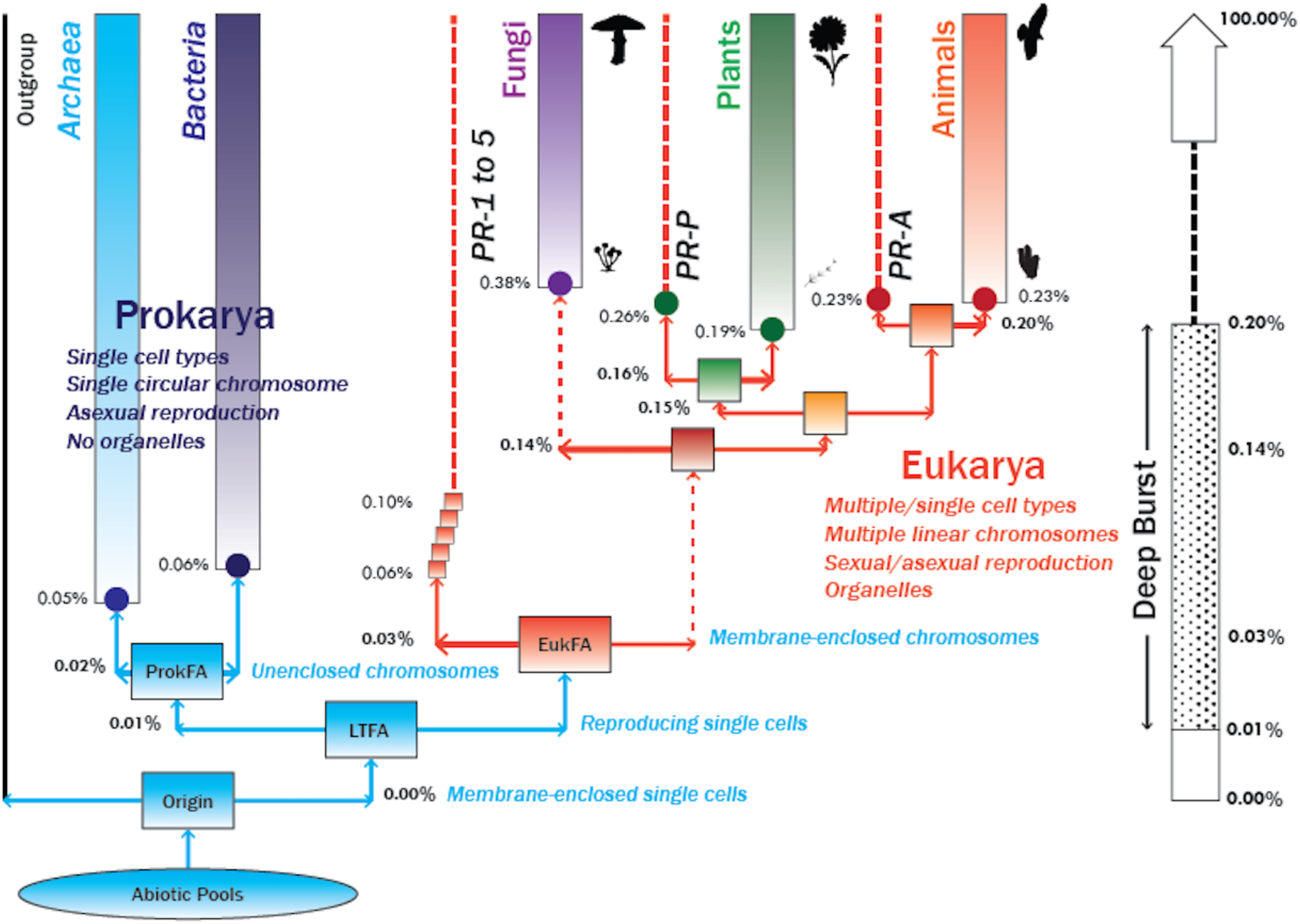
Simplified genome ToL at the deepest level. All extant organisms in this study are grouped into five “Majorgroups”, as shown as five columns (corresponding to Archaea, Bacteria, Fungi, Plants and Animals) and a paraphyletic protist “Group” represented by three thin dotted red columns (corresponding to three groups of protists, labeled as PR- 1~5, PR-P, and PR-A). For simplicity, “singletons”, the organisms that do not belong to any named groups, are not shown. It also shows the cumulative branch-lengths to internal nodes (rectangles). Each small circle represents the internal node of the clade containing all *extant* members of the clade of a Majorgroup, PR-P or PR-A subgroup, and each rectangular box presents a pool of the founding ancestors (FAs) from which one or more founders *emerge* (or are selected) to evolve to become a node containing an *extant* organism or the “seed” for the next founder pool (see Supplemental Fig. S2). The bold number next to each horizontal arrow is the Cumulative Genomic Divergence (CGD) value (cumulative branch-length), at which the founder(s) of the respective Majorgroup emerged, and the plain number next to each circle corresponds to the CGD value of the internal node of a clade containing all the extant members of the Majorgroup. The silhouettes of one of the early emerged and one of late emerged organisms of each Majorgroup among the study population are shown: a small member of Ascomycota and a mushroom for Fungi, an algae and a flowering land plant for Plants and a sponge and a bird for Animals, respectively. LTFA: the pool of Last Terrestrial (Earth-bound) Founding Ancestor of replicating cells; Origin: the last pool of non-replicating cells of diverse contents, from which the founders of LTFA emerged; ProkFA: the pool of Prokarya founding ancestors; EukFA: the pool of Eukarya founding ancestors.

The membership of each of all eukaryotic Majorgroups and Groups in the genome ToL coincide with those of the groups identified by NCBI taxonomic names at a Phylum (P), a Class (C) or an Order (O) level with a few exceptions marked by asterisks before the taxonomic names (see Fig. 1 legend). A notable exception are the three Groups of Majorgroup Protists (see Three groups of Majorgroup Protists in E. Phylogeny of Supergroup Eukarya below).

For prokaryotes, this is also the case at Major group level, but not at Group level (see Phylogeny of Supergroup Prokarya (or Akarya) below)). However, as described below, there are significant differences in the topological relationship among the groups, relative branch-lengths, and branching order associated with the groups between the genome ToL and geneToLs (see below).

### B. First emergence of the “founders” of Supergroup Prokarya and Supergroup Eukarya

Figure 2 shows that the *founder* (see Pool of “founding ancestors” vs. common ancestor in Introduction above and Supplement Fig. S2) of the Supergroup Prokarya and the founder of the Supergroup Eukarya emerge first (“Eukarya early” model) from the pool of Last “Terrestrial (Earth-bound)” Founding Ancestors (LTFA; analogous to LUCA (Last Universal Common Ancestor), but emphasizing the aspects of *terrestrial* ancestors and a *pool* of founding ancestors of the first internal node of the genome ToL). Then, the founders of Majorgroups Archaea and Bacteria and those of all eukaryotic Majorgroups (Protists, Fungi, Plants and Animals) emerge from their respective Supergroup founding ancestors (FAs) pools. This is in contrast to the branching out of Archaea and Bacteria from LUCA first, followed by branching out of Eukarya from Archaea accompanied by one or more steps of fusion events between some members of bacteria and archaea (“Eukarya late” model), according to most of the recent gene ToLs (4, 6, 7, 8, 9, 23, 24). However, it is in agreement with our earlier rooted genome tree of prokaryotes constructed using the proteome sequences of 884 prokaryotes plus 16 unicellular eukaryotes (17), and a ToL reconstructed using the contents of coding sequences for protein “fold-domain superfamily” and rooted in a way different from most gene ToLs and our genome ToL (22).

### C. “Deep Burst” of the founders of all Majorgroups

Figure 2 shows Cumulative Genomic Divergence (CGD) values, which are the cumulative branch-lengths along the lineage for all the founding ancestor nodes (internal nodes) of the 5 Majorgroups and 3 protist Groups. CGD values are scaled from 0 % for the Origin of Life to 100% for the most recent extant organisms sequenced (see Chronological time scale vs. Progression scale in Discussion). The figure also highlights that the founders for all Majorgroups in the study population have *emerged abruptly* during a three-staged “Deep Burst” of genomic divergence near the very beginning of the Origin: (a) At the first stage (CGD scale of 0.01%) the founding ancestors of the two Supergroups, Prokarya and Eukarya, *emerge* from LTFA; (b) In the second stage (CGD scale of 0.02 - 0.03%), the founding ancestors of three types of *unicellular* organism groups emerge: Archaea and Bacteria emerge from the Prokarya pool of founding ancestors, and the founders of the unicellular protist Groups (PR-1 to 5) emerge from the Eukarya pool of founding ancestors; and (c) In the third stage, (CGD of 0.14 - 0.20%), the founders of three Majorgroups corresponding to Fungus, Plant and Animal clades plus two protist groups (PR-P and PR-A) emerge. Thus, the abrupt emergence of the founders of all five Majorgroups plus a Protist Majorgroup, composed of three types of protists, occurred during the Deep Burst within a very short range between 0.01% and 0.20% on the *progression* (*i.e. CGD*) *scale of evolution*, followed by 99.8% of CGD of “gradual” evolution of the Majorgroups (see Chronological time scale vs. Progression scale in Discussions).

### D. Emergence and Divergence of the Founders of 35+ Groups

Figures 3 and 4 show that each founder of 35+ Groups emerged during a period corresponding to CGD scale between 0.05% (emergence of the founder of Archaea in Fig. 2) and 27.62% (emergence of the founders of extant birds and crocodile/turtle Groups in Fig. 3), suggesting that, depending on the Group, 99.95% to 72.38% in CGD scale account for the “relatively gradual” (not “bursting”) evolution toward the extant organisms Groups.

**Fig. 3.**
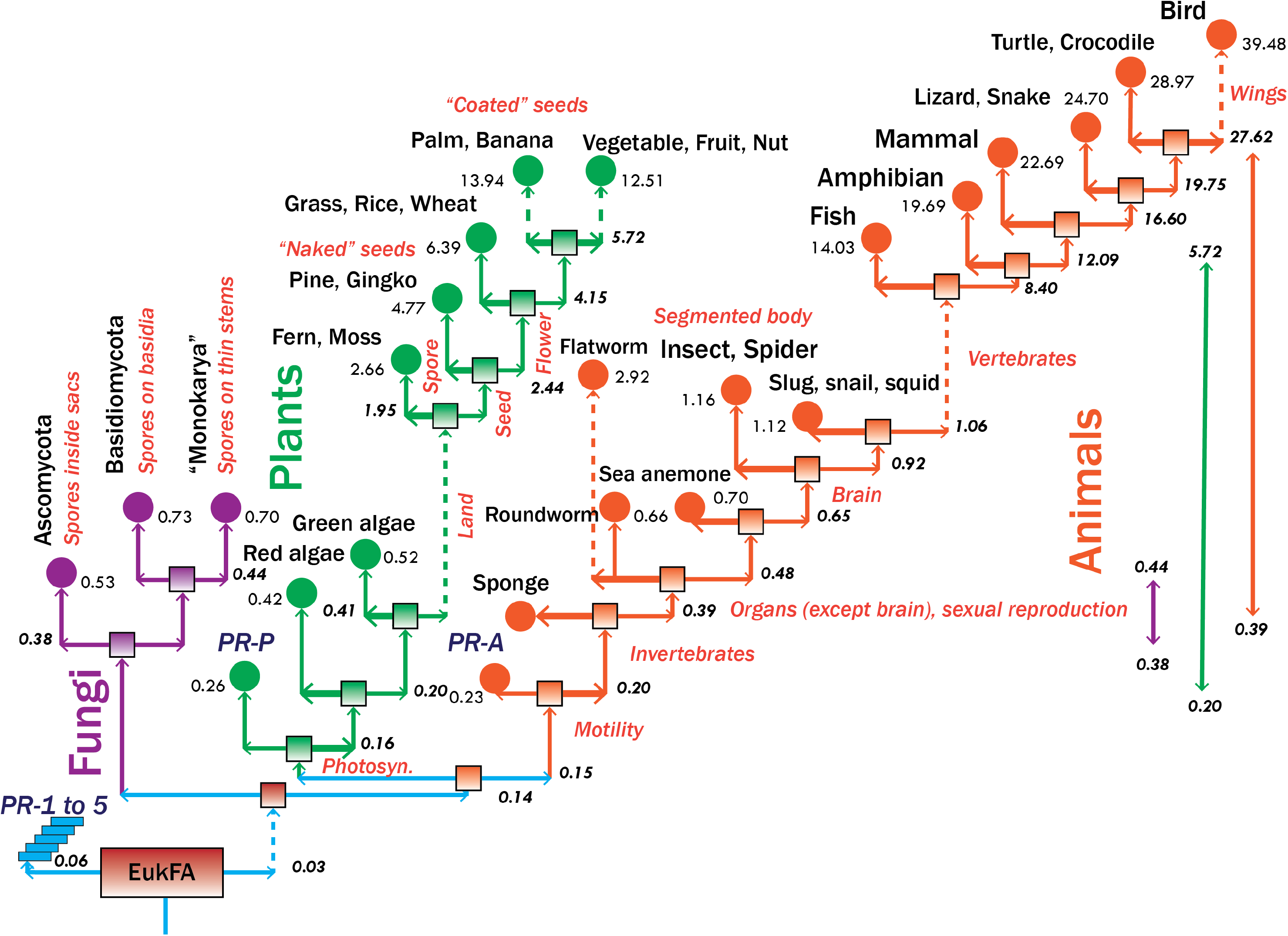
Simplified eukaryotic portion of the genome ToL at Group level. The portion of the genome ToL corresponding to all eukaryotes are shown at the tree-branching level of Groups. For simplicity, “singletons” (that do not belong to any named groups). Also not shown are all leaf nodes and their branches. Rectangles and circles as well as the bold number next to each horizontal arrow and the plain number next to each circle have the same meanings as those in Fig. 2. The dotted vertical arrows are to indicate that they are arbitrarily shortened to accommodate large jumps of CGD values within a limited space of the figure. The percentage symbols of CGDs are not shown. The range of the CGD values within which all the founders of the extant organisms of each Majorgroup have emerged are shown on the right. The CGD values of each colored Majorgroup are not scaled to those of the others.

**Fig. 4:**
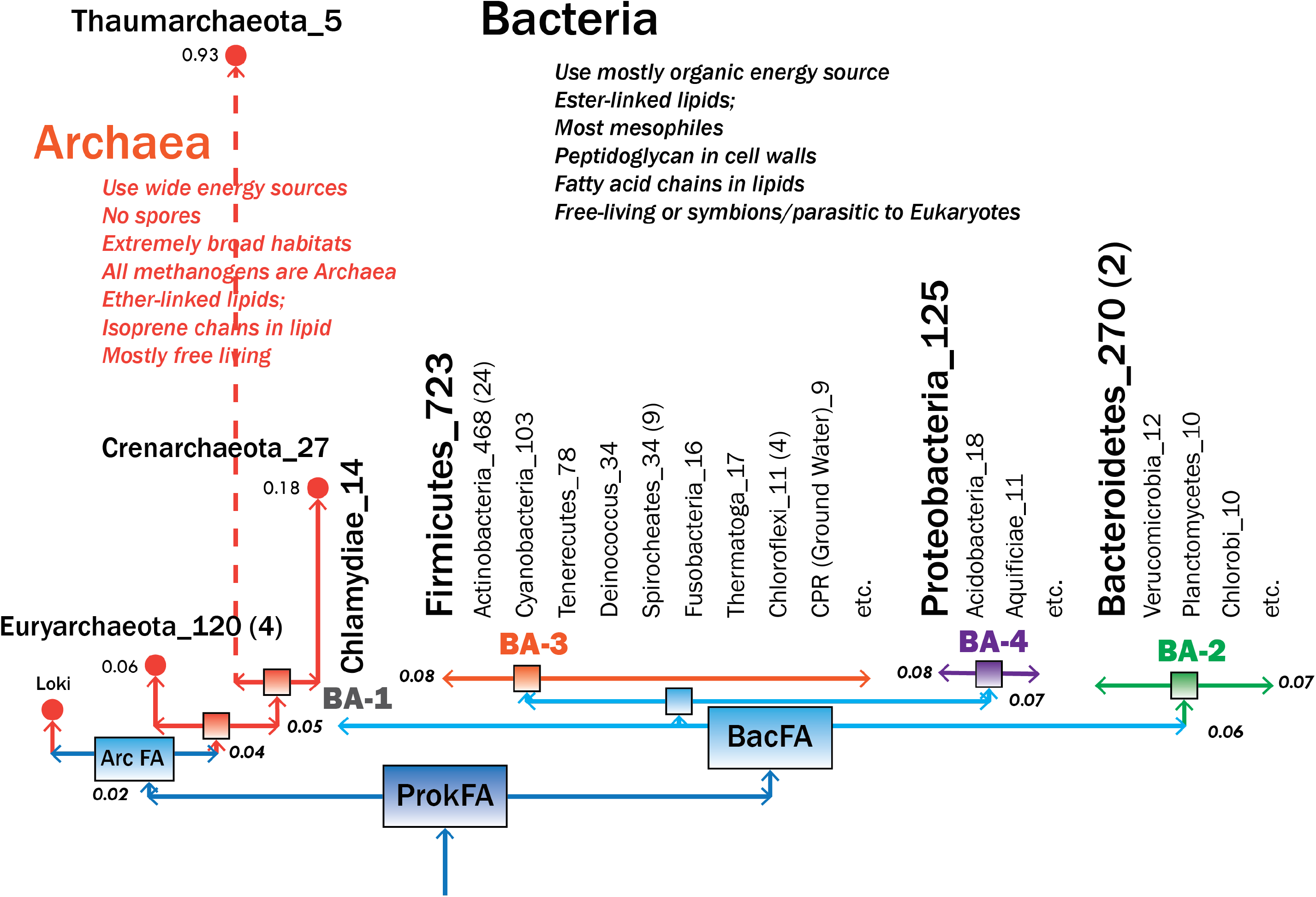
Simplified prokaryotic portion of the genome ToL at Group level. The portion of the genome ToL for prokaryotes are shown at Group (mostly phylum), where, for simplicity, Subgroups with small sample sizes and “singletons” are not shown. Majorgroup Bacteria is divided into four Groups, BA-1 to 4, where each Group consists of one or many Subgroups. The bold faced name in each Group has the largest sample size among the members of the Group. Number after each NCBI taxonomic name refers to the sample size of the majority clade, followed by a number in parenthesis referring to the size of minority clade away from the majority clade. Interestingly, the founders of all the named Groups of Bacteria emerged within a very small range of CGD values of 0.06% - 0.08%, in a drastic contrast to a much larger range of 0.20% - 27.62% for those of all eukaryotes (The numbers next to horizontal lines and circles have the same meanings as those in Fig. 2.). Thus, the order of the emergence of all the named bacterial Groups and Subgroups may be less reliable compared to that of eukaryotes. For simplicity, the branching order among the Subgroups is not shown.

### E. Phylogeny of Supergroup Eukarya

#### Branching orders within three Majorgroups

Figure 3 shows the order of the emergence of the founders of all eukaryotic Majorgroups. The branching order of the three Majorgroups (Fungi, Plants, and Animals) differs from those of the gene ToLs: Almost all recent gene ToLs show Fungi as the sister clade of Animal clade (for a recent review, see 23), but in our genome ToL, Majorgroup Fungi is sister to the combined group of the Plant *and* Animal Majorgroups plus their respective Protist sister Groups, PR-P and PR-A..

For Majorgroup Fungi, as reported earlier (18), the founders of all three Groups of Fungi corresponding to Ascomycota, Basidiomycota and “Monokarya” (“Non-dikarya”) emerged within a small CGD range of 0.38 – 0.44 %, of which the founders of the Ascomycota appears first at CGD of 0.38%, around the similar value of CDG when the founders of red and green algae of Plants and of invertebrates of Animals emerged.

In Majorgroup Plants, the order of emergence of the founders starts with those of marine plants, such as red algae and green algae. After a large jump of CGD the founders of non-flowering land plants such as spore-forming ferns and land mosses emerged, then “naked” seed-forming Gymnosperms such as gingko and pines, followed by “enclosed” seed-forming flowering land plants of Angiosperms such as Monocot and Dicot plants.

In Majorgroup Animals, the order of the emergence of the founders starts with those of invertebrates such as sponges, worms, cnidaria, arthropods, and mollusca. Then, after a big jump of CGD, the founders of vertebrates such as fishes, amphibians, mammals, and reptile-A (snakes, lizards, etc.), emerge, and finally, the sister pair of the founders of reptile-B (crocodiles, alligators etc.) and birds, of which the extant birds with wings emerge after another big jump of CGD.

There are 22 Groups among all three Majorgroups in this study population, and all founders of the Groups emerged in a very wide range of CGD between 0.20% and 27.48% (see Fig. 3). This is in drastic contrast to the very small CGD range of 0.06% and 0.12% in which all founders of Prokaryotic Groups emerged (see Phylogeny of Supergroup Prokarya below).

#### Three types of Majorgroup Protists

Protists are currently defined as unicellular eukaryotic organisms that are not fungi, plants or animals. Despite the paucity of the whole genome information of protists, especially of a large population of non-parasitic protists, the genome ToL suggests that there are at least 7 subgroups of protists in three types as mentioned earlier. Figure 2 shows that the protist subgroup PR-P is the sister clade to Majorgroup Plants (red and green algae and land plants), and most of them belong to one of the two categories: photosynthetic protists, such as microalgae, diatoms, phytoplanktons, or water molds parasitic to plants. Another protist clade, PR-A, is a larger subgroup including many protists with a wide range of phenotypes and is the sister clade to Majorgroup Animals. Most of them are motile and can be grouped with slime molds, amoeboids, choanoflagellates, and others, or with parasites to cattle and poultry (see Fig. 3). The rest of protists form 5 small subgroups (labeled as PR-1 to PR-5 in Figs. 1, 2 and 3), which do not form a larger clade together, but emerge successively from the pool of Eukarya founding ancestor. Most of them are parasitic to animals with varying host specificities. The founders of these protist subgroups emerged much earlier than those of PR-P and PR-A as implied by smaller CGDs, and they correspond to Entamoeba, Microsporidia, Kinetoplastida, Oligohymenophorea and Apicomplexa, respectively.

### F. Phylogeny of Supergroup Prokarya (or Akarya)

Deep branching pattern of Supergroup Prokarya is much more “collapsed” beyond the Majorgroup level compared to that in Supergroup Eukarya. Figure 4 shows a simplified prokaryotic portion of the genome ToL. At the deepest level of Supergroup Prokarya (CGD of 0.02% from the Origin), the founders of two Majorgroups, corresponding to Archaea and Bacteria, emerge, followed by the emergence of the founders of three Groups of Archaea (AR-1, 2, and 3 corresponding approximately to the extant Euryarchaeota, Thaumarchaeota and Crenarchaeota) and four large bacterial Groups (BA-1 to BA-4), with no obvious distinguishing characteristics, within a very small CGD range of 0.05 – 0.06% (Fig. 4; see also On the Phylogeny of Supergroup Prokarya (or Akarya) in Discussion). Of the four Groups, BA-1 has only one member, Chlamydia, obligate intracellular parasites, and is basal to the remaining three. BA-3 is the largest and is very similar to an unranked “Terrabacteria super group”, whose habitat is strictly “non-marine” (e.g., soil or rock on land; freshwater of lakes, rivers, and springs), or if their host is a non-marine species (25). Members of this Group include those resistant to environmental hazards such as ultraviolet radiation, desiccation and high salinity, as well as those that do oxygenic photosynthesis. The remaining two Groups have not been named in the NCBI database.

Among the four bacterial Groups in this study population, there are more than 30 bacterial Subgroups corresponding to the groups with NCBI taxon names at the Phylum level. The founders of all of them emerged within a small CGD range of 0.06% and 0.12% (Fig. 4), thus, making it less certain about the resolution of the branching order not only among the four Groups but also among many Subgroups within each Group. This is a drastic contrast to a very large CGD range between 0.20% and 27.48% in which all founders of Eukaryotic Groups emerged (see Phylogeny of Supergroup Eukarya above).

### G. Phylogenic positions of new groups

Candidate Phyla Radiation (CPR) group is a newly discovered very large group of small bacteria (estimated to be about 15% of all bacteria) with relatively small genomes. Most of them are found in diverse environments including ground water, and are symbiotic with other microbes in their community, thus, difficult to culture their representative members for genome sequencing. An extensive metagenomic studies and gene tree construction using 16 ribosomal protein genes revealed that the CPR group members form a “super group”, which is well-separated from (or a sister to) *all* other Bacteria (7). Despite its vast size, only 8 genome sequences are available at present in public databases. In our genome ToL these small samples form a sister clade to Tenericutes, a member of Majorgroup BA-3 of Bacteria (see Fig. 4). More full genome sequences of other CPR group members may resolve this apparent discrepancy of clading and interpretation.

Hemimastigophora group is one of the unicellular eukaryotic protist groups with uncertain phylogenic assignment due to the absence of genomic sequence information. The members of the group are mostly free-living and predatory to other microbes in soil, sediment, water column and soft water environment, and have highly distinctive morphology. Based on recent studies of transcriptome sequences of two members of the group, an un-rooted gene tree using 351 common genes was constructed and reported that Hemimastigophoras form a new “supra-kingdom” at the basal position of the Animal kingdom (26). Our genome ToL is showing the group as a member of the protist subgroup of PR-A, which is the basal or sister clade to Majorgroup Animals.

### H. Phylogenic positions of “singletons”

Figure 1 and Supplemental Fig. S3 show the complete genome ToL grouped at phylum, class or order level: the former as a circular topological ToL ignoring the branch-lengths, and the latter as a linear ToL with all the cumulative branch-lengths shown for internal nodes from which all the founders of the groups emerge. In both figures most of “singletons” are included. “Singleton” is defined, for this study population, as an organism that does not find a closest neighbor of the same group name. Many of them are at basal/sister positions to larger groups suggesting possible *speculative* evolutionary roles, accompanied with “accumulative” or/and “reductive” evolution, and one suggests reassignment of group affiliation:

(a). Lokiarchaeum sp as a basal organism to all members of Majorgroup Archaea, rather than a member of the sister group to Euryarchaeota (23)

(b). *Giardia lamblia* and *Trichomonas vaginalis* at the basal or sister position to Supergroup Eukarya.

(c). *Naegleria gruberi* at the basal position of three Majorgroups of Fungi, Plants and Animals plus two protist clades, PR-P and PR-A.

(d) *Emiliania huxleyi* and *Guillardia theta* as a nearest neighbor pair at the basal position of Majorgroup Plants.

(e). *Thecamonas trahens* at the basal position of Majorgroup Animals plus the protist group PR-A.

(f). *Sphaeroforma arctica* and *Amphimedon queensiandica* at the basal position of Majorgroup Animals (27)

(g). *Trichoplax adhaerens* at the basal/sister position to Cnidaria clade (slug, snail, squid) of Majorgroup Animals.

(h). *Ciona intestinalis* and *Hellopdella robusta* emerging between Cnidaria clade and Arthropoda clade (insect, spider) of Majorgroup Animals.

(i). *Branchiostomas* emerging before Group Fish of Majorgroup Animals.

(j). *Callorhinchus milli* and *Latimeria chalumnae* emerging between Group Fish and Group Amphibian of Majorgroup Animals.

The genome sequences of more organisms that are close relatives of these singletons are needed to confirm or refute these speculations.

## Discussions

### Chronological time scale vs. Progression scale

There is no known measure to estimate the *chronological evolutionary* time along the evolutionary lineage of an extant organism, especially in deep evolutionary period. However, the degree of *evolutionary progression* from the Origin (0% progression) to the extant organism (100% progression) can be scaled to the minimum and maximum of JSD, which is bound between 0 and 1. Thus, the % progression of evolution from the Origin to a given internal node can be represented by the cumulative branch length (CGD value) of the internal node. Although the chronological evolutionary time scale is different from the evolutionary progression scale, both scales have the *same directional arrow* starting from the Origin to the most recent extant organisms, thus, the *ranking order* (*branching order*) of the emergence of the founders along the *lineage will be the same*.

### “Deep Burst” model vs. other models for evolution of organisms

As mentioned earlier in Results, the founders of all Majorgroups emerged by the time corresponding to 0.20% on CGD scale (see Fig. 2). Thus, the remaining 99.80% of CGD scale accounts for presumed multiple “relatively” gradual (less abrupt) evolutionary steps, multiple cycles of emergence of new founders and gradual divergence of their genomes. Such explosive birth model of Deep Burst has notable similarities to various aspects of earlier models of evolution of Life: (a) The non-tree like “Biological Big Bang (BBB)” hypothesis (28), especially the second of the three BBBs, is similar to the emergence of the first stage of the Deep Burst and (b) the un-resolved “Bush-like tree” model (29), especially the “collapsed” aspect of deep branches at the Group level of Prokarya. Both models are inferred from various analyses of gene ToLs and they appear to correspond to one of the three stages of the Deep Burst, all occurred during the period corresponding to 0.20% on CGD scale. The remaining 99.8% of CGD scale accounts for the gradual evolutionary steps, after the Deep Burst of emergence of all the founders of Majorgroup, has similarity to “Punctuated Equilibrium” model (30) inferred from paleontological analyses of fossils.

### On the phylogeny of Group Animal

The sequence of the emergence of the *founders* of all named Groups as shown in Fig. 3 agrees with that of most of the gene ToLs as well as fossil data except that of mammals and birds. Many gene ToLs and fossil data suggest mammals emerging *after* emergence of birds and reptiles. In our genome ToL, the *founders* of Group Mammal is sister to those of the joined Groups of birds and two types of reptiles. Such sisterhood was also detected in some gene ToLs (e.g., 4). Furthermore, Fig. 3 also shows that the founding ancestor of the *extant* mammal Group of this study population, indicated as circles in Fig. 3, emerged earlier than both of the *extant* bird Group and the *extant* reptile Groups. This is “counter-intuitive”, although one can imagine a narrative that all pre-mammal birds and reptiles did not survive certain mass-extinction event(s), but some of those are detected only as fossils.

### On the Phylogeny of Supergroup Prokarya (or Akarya)

For all eukaryotic organisms in the study, the membership of the organisms in each of the 21 Groups (3 fungus Groups, 7 plant Groups, and 11 animal Groups; 3 protist Groups are not counted) agrees well with that of the organisms in most gene ToLs at phylum/class names (see Fig. 3). This is also true for most prokaryotic groups (about 30 groups at phylum level) with some exceptions, where one or more small minority of a group does not clade with their respective majority group (see Fig. 4). Some of many possible narratives for these minority “discrepancies” are: (a) Among prokaryotes, extensive genomic transfer may have occurred “horizontally” early in their evolution such that some of the boundaries among various groups became not clearly resolvable at a whole genome level, (b) Assigning a name of a new prokaryotic organism based almost exclusively on the clading pattern in gene trees, constructed from a very small number of *selected genes*, may not be identical from that in genome ToL, constructed from almost *all genes* of the organism, (c) Deep branching pattern of Supergroup Prokarya is highly “collapsed” (fig. 4), thus, may be less reliable beyond Majorgroup level, and/or (d) misclassification at the time of naming, or some unknown “artifacts” of clading algorithm between two different “distance measure” used. These discrepancies are expected to be resolved once more whole genome sequences of diverse members of the small minorities become available.

### Similar composition of each clade but different branching order of the clades

We found it surprising that, although the *branching order* of various groups are significantly different between gene ToLs and our genome Tol, the *membership* of each group of *almost all Groups* (29 out of 32 non-Protist Groups) is the *same not* only at all Supergroup and Majorgroup levels, but also at the 29 Group level (corresponding to phylum). This is especially surprising because the membership of each clade is assigned by the clading pattern, which is determined by two completely different ways. One possible explanation is that, this surprising observation is the consequence that, after the Deep Burst, when the founders of all groups emerged, all the organisms within each group evolved “isolated” from other groups for the evolutionary time corresponding to most of CGD scale (see Fig. 2). We attribute the differences in the branching order of the groups to the fact that the branch lengths are calculated by two totally different distance measures applied on two completely different descriptors for the organisms.

### A narrative

Figure 2 is a simplified genome ToL, which suggests possible narratives of the evolutionary course and kinship among the Majorgroups of extant organisms in this study. It also suggests that the founders of all the Majorgroups emerged in a Deep Burst, almost at the time of the start of Life on Earth from a population of cells of diverse size and contents (LTFA) which may have been formed by random packaging of various assortments of molecules in an aqueous pool under a critical environmental condition.

This narrative is contrasted with those from most current gene ToLs in Supplementary Fig. S4, which emphasizes the differences in grouping, branching order of the groups, and the nature of the internal nodes between the two types of ToLs.

## Materials and Methods

### Sources and selection of proteome sequences and taxonomic names

All publicly available proteome sequences used in this study are obtained from National Center for Biotechnology Information (NCBI). We downloaded the proteome sequences for 691 eukaryotes and 3317 prokaryotes from NCBI RefSeq DB using NCBI FTP depository (32; as of July, 2017, our project start time). Proteome sequences derived from all organelles were excluded from this study. In addition, we included the proteome sequences of 9 prokaryotes (as of August 2017): Lokiarchaeum (23) and 8 CPR ground water bacteria (7) from NCBI genebank, and 4 gymnosperms (*Ginkgo biloba*, *Pinus Lambertiana*, *Pinus Taeda* and *Pseudotuga Menziesi*) from Treegenesdb (33; as of Jun 2018). Thus, the total of 4023 proteome sequences form the population of this study.

Proteome sequences not included in our study are those derived from whole genome sequences assembled with “low” completeness based on two criteria: (a) the genome assembly level indicated by NCBI as “contig” or lower (i.e. we *selected* those with the assembly levels of ‘scaffold’, ‘chromosome’ or ‘complete genome’), and (b) the proteome size smaller than the smallest proteome size among highly assembled genomes of eukaryotes and prokaryotes, respectively. For the minimum proteome size threshold for eukaryotes, we used 1831 protein sequences of *Encephalitozoon romaleae* SJ-2008 (TAXID: 1178016) and that for prokaryotes we used 253 protein sequences from *Candidatus Portiera aleyrodidarum* BT-B-HRs (TAXID: 1206109).

All taxonomic names and their taxon identifiers (TAXIDs) are based on NCBI taxonomy database (34).

### Feature Frequency Profile (FFP) method

The method (14) and two examples of the application of the method (17, 18) have been published. A brief summary of the two steps taken specifically for this study is described below:

In the first step, we describe the proteome sequence of each organism by the collection of all unique n-grams (Features), which are short peptide fragments, generated by a sliding “window” of 13 amino-acids wide along the whole proteome sequence of the organism. Some Features may be present more than once, so we log the counts. This n-ram contains the *complete information* to reconstruct the whole proteome sequence of the organism. Then, since each organism’s proteome has a different size, we convert all the counts of Features to frequencies by dividing by the total number of counts for each proteome. Thus, now, each organism is represented by the FFP of its proteome sequence.

The second step is comparing the two FFPs to measure the degree of difference ("divergence") between the two FFPs by Jensen-Shannon Divergence (JSD; see next section below), which measures the degree of difference between two proteome sequences, that is, two FFPs. All pairwise JSDs, then, form a "divergence distance matrix", from which we construct the genome ToL and all the branch lengths (see below).

### Jensen-Shannon Divergence (JSD) and cumulative branch-length

JSD values are bound between 0 and 1, and they correspond to the JSD value between two FFPs of identical proteome sequences and two completely different proteome sequences, respectively. Any differences caused by point substitutions, indels, inversion, recombination, loss/gain of genes, etc., as well as other unknown mechanisms, will bring JSD somewhere between 0.0 and 1.0 depending on the degree of *information divergence*. In this study the collection of the JSDs for all pairs of extant organisms plus 4 out-group members constitute the divergence distance matrix. BIONJ (35, 36) is used to construct the genome ToL. For convenience of comparison, all branch-lengths are scaled to produce the cumulative branch length of 100% from the Origin to the most recent extant organism. (In practice, this is done by scaling the maximum value of JSD to 200%.) A cumulative genomic divergence (CGD) of an internal node is defined as the cumulative sum of all the scaled branch-lengths from the Origin to the node along the presumed evolutionary lineage of the node.

### Choice of the “best” descriptor and “optimal” Feature length

In this study, two key decisions to be made are the choice of the *descriptor* for the whole genome information system (DNA sequence of genome, RNA sequence of transcriptome or amino acid sequence of proteome) and the choice of the optimal Feature length of FFP to calculate the *divergence distance* between a pair of FFPs. Since there is no *a priori* criteria to guide the making of the choices (for other choices, see 37) we took an empirical approach, learned from our earlier studies of building whole genome trees for the kingdoms of prokaryotes and fungi (17, 18), where we took an operational criterion that the “best” choice should produce the *most stable tree topology*, as measured by Robinson-Foulds (R-F) metric (38) in PHYLIP package (39), which estimates the topological difference between two ToLs, one with optimal Feature length of l and the other with l+1. The results of the search showed (supplemental Fig. S1) that, among the three types of genome ToLs, the proteome-sequence based genome ToL is most topologically stable because it converges to the ToL with lowest R-F metric and remains so for largest range of Feature length starting from Feature length of about 12. As for the physical meaning of the optimal Feature length, we can infer it from the experiment with books without spaces and delimiters (14), where it approximately corresponds to the Feature length at which the number of “vocabulary”, the Features with unique sequences, is the maxium among all books compared (14).

### “Out-group” members

For the out-group of our study we used the *shuffled proteome sequences*(40, 41) of two eukaryotic and 2 prokaryotic organisms: For prokaryoyes we chose *Candidatus Portiera aleyrodidarum* BT-B-Hrs (Gram-negative proteobacteria) with the smallest proteome size of 253 proteins and *Ktedonobacter racemifer* DSM 44,963 (green nonsulfur bacteria) with the largest proteome size of 11,288 proteins; For eukayotes we chose two fungi: a Microsporidia, *E. romaleae* SJ-2008, with the smallest proteome size of 1,831 proteins and a Basidiomycota, *Sphaerobolus stellatus*, with the largest proteome size of 35,274 proteins.

## Supporting information

ToL.PNAS.Suppl Info

## Computer code availability

The FFP programs for this study (2v.2.1) written in GCC(g++) is available in Github: https://github.com/jaejinchoi/FFP.

## Web address links;

TransDecoder for translating the transcriptome sequence to amino acid sequence: https://github.com/TransDecoder/TransDecoder/wiki treegenesdb: https://treegenesdb.org/, FTP: https://treegenesdb.org/FTP/

Hemimastigotes transcriptome data from https://datadryad.org/resource/doi:10.5061/dryad.n5g39d7

### Acknowledgement

We thank Dr. Byung-Ju Kim (formerly at University California, Berkeley, CA, USA, Yonsei University, Seoul, Korea, and currently at Incheon National University, Incheon, Korea) for numerous discussions and advice on computer programming and algorithm development relevant to this study. We also gratefully acknowledge the expert comments and advices from our colleagues at University of California, Berkeley, CA.: Prof. John Taylor on Fungi, Profs. James Patton and David Wake on mammals and reptiles, Profs. Alex Glaser and Hiroshi Nikaido on Prokarya, Profs. Rauri Bowie on Birds, Profs. Peter Oboyski and Kip Will on Insects, Prof. Bruce Baldwin on Plants and Prof. Nicole King on Protists. Special thanks to Prof. Se-Ran Jun of College of Medicine, University of Arkansas for Medical Sciences, Little Rock, AR and Prof. Norman Pace of University of Colorado, Boulder CO for their constructive comments and advice on our first draft of the paper.

This research was partly supported by a grant (to SHK) from World Class University Project, Ministry of Education, Science and Technology, Republic of Korea and a gift grant to University of California, Berkeley, CA. (to support JJC).

SHK acknowledges having an appointment as Visiting Professorships at Yonsei University and Incheon National University in South Korea during the manuscript preparation.

## Author contribution

Conceptual design of the study and speculative interpretations of the results by SHK; downloading and curation of sequence data from various data bases, computational algorithm design, and programming and execution by JJC; interpretation of computational results by SHK and JJC; manuscript preparation by SHK in extensive consultation with JJC; all figures are designed by JJC and SHK.

