## Supplementary material for "Genome Tree of Life: *Deep Burst* of Organism Diversity": ToL.PNAS.Suppl Info

**Supplementary Information**

**Legends for Supplementary Figures**

**Figure S1. Optimization of tree topology as a function of Feature length.**

Topological variation, as represented by Robinson-Foulds metric (38) between a pair of trees, is shown on Y-axis, and Feature length on X-axis. Each tree is constructed using a “divergence distance matrix” calculated by Jensen-Shannon divergence (JSD; 16) for all pair-wise Feature Frequency Profiles (FFPs). Robinson-Foulds metric is calculated between two trees: one for Feature length of l and the other for Feature length of l +1. This experiment is done by sampling 301 from 4023 species used in this study for computational expediency. This figure shows that, among three types of genomic information (DNA sequence of whole genome, RNA sequence of transcriptome, and amino acid sequence of proteome), the proteome sequence-based ToL converges to the most topologically stable tree starting from the Feature length of 12 (having the lowest point in the curve) and remain stable for longer Feature lengths. For this study we use Feature length of 13.

**Figure S2. Internal node as a Pool of founding ancestors**

An internal node (shown as a rectangle) is represented as “a pool of diverse founding ancestors (FA)” like a pool of mosaic tiles. Each internal branch is divided into two components: the horizontal arrows to represent the emergence of one or more *founders* from the pool under “abrupt” local environmental and ecological changes, and the vertical arrows to represent the genomic *diversification* of the founders to evolve into a new pool of founding ancestors through relatively gradual evolution. An internal node of a clade containing all extant members of a named group (such as bacteria, plants, fish, etc.) is shown as a circle

**Figure S3. Genome ToL with scaled cumulative branch-lengths**

The genome ToL consists of all study organisms (697 eukaryotes, 3326 prokaryotes, and 4 outgroup members) grouped at Phylum, Class, or Order level. Common or scientific names for the clading groups are shown, followed by sample numbers as in Fig. 1. “Singletons” are indicated by their scientific names. Coloring scheme is the same as that in Fig.1. The scaled cumulative branch-length (see Jensen-Shannon Divergence and Cumulative Genomic Divergence in Materials and Methods) corresponding to CGD larger than 1.0% are proportionately shortened and seen as dotted-line while maintaining its ranking order. The computer visualization of ToL was made using ITOL(42). Coloring and labeling schemes on left are the same as those of Fig. 1.

**Figure S4.** Schematic representation of the genome ToL (left) and one version of current gene ToLs (right). In the genome ToL, two Supergroups (Akarya (Prokarya) and Eukarya) are colored purple and all Majorgroups ( Archaea, Bacteria, Fungi, Plants, Animals, and three Protists (dotted lines)) orange, and all the internal nodes are shown as circles of variable size to emphasize that each one is a *pool* of founding ancestors with *wide genomic divergence*, like a bucket of different mosaic tiles. In the gene ToL, three Domains are shown in different colors (Bacteria in green, Archaea in blue, and Eukarya in red) and the internal nodes are shown as the circles of same size to imply that each node consists of one or more clones with high genomic homology, i.e., *little genomic divergence*. The dotted arrow suggests one or more steps of cell fusion processes between some members of Bacteria and Archaea.

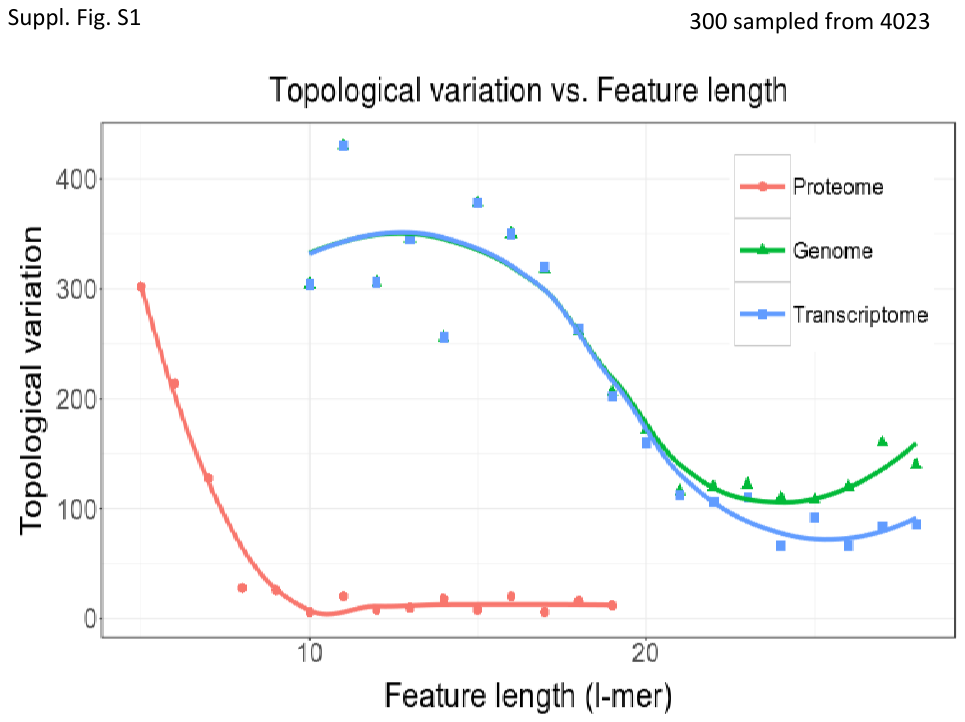

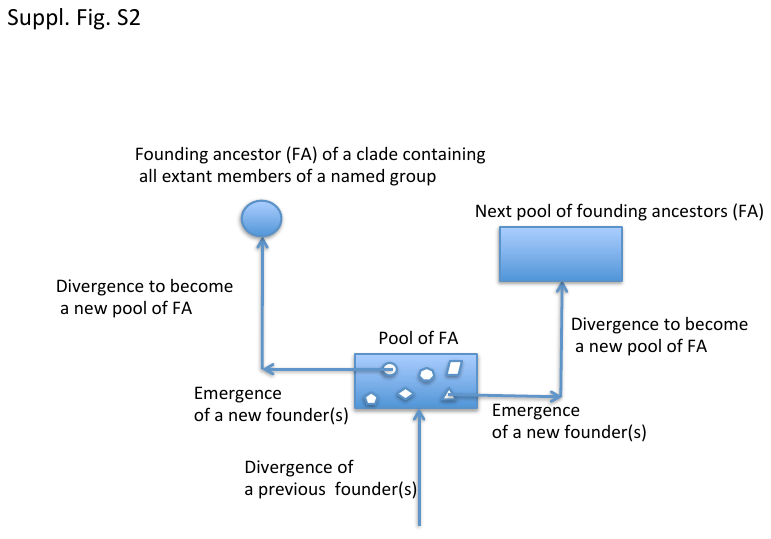

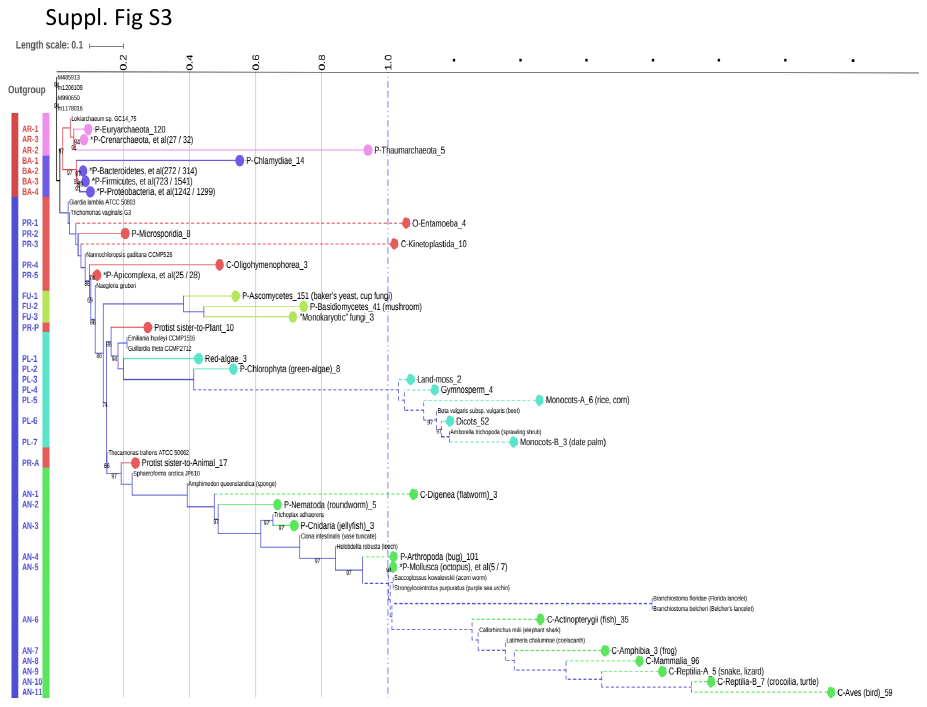

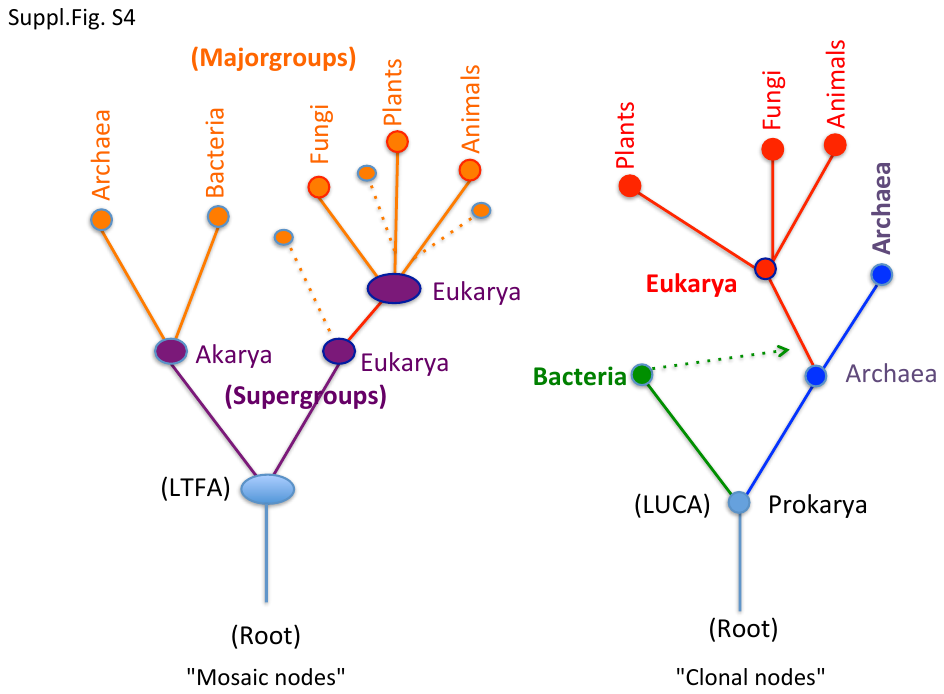
